# Single cell in vivo brain optogenetic stimulation by two-photon excitation fluorescence transfer

**DOI:** 10.1101/2020.06.29.179077

**Authors:** Lei Tong, Peng Yuan, Yao Xue, Minggang Chen, Fuyi Chen, Joerg Bewersdorf, Z. Jimmy Zhou, Jaime Grutzendler

## Abstract

Optogenetic manipulation with single-cell resolution can be achieved by two-photon excitation; however, this frequently requires relatively high laser powers or holographic illumination. Here we developed a practical strategy to improve the efficiency of two-photon stimulation by positioning fluorescent proteins or small fluorescent molecules with high two-photon cross-sections in the vicinity of opsins. This generates a highly localized source of endogenous single-photon illumination that can be tailored to match the optimal opsin absorbance. Through neuronal and vascular stimulation in the live mouse brain, we demonstrate the utility of this technique to achieve more efficient opsin stimulation, without loss of cellular resolution. We also provide a theoretical framework for understanding the potential advantages and constrains of this methodology, with suggestions for future improvements. Altogether, this fluorescence transfer illumination method allows experiments difficult to implement in the live brain such as all-optical neural interrogation and control of regional cerebral blood flow.

## Introduction

Optogenetics with light-sensitive opsins has revolutionized the field of neuroscience (1). Typical experiments involve the utilization of single-photon illumination of the brain to activate opsins such as channelrhodopsin (ChR2) and others (2). While this allows temporally precise manipulation of cell ensembles, all the cells along the conical illumination light path are activated, reducing the spatial specificity and resulting in artificially synchronized activity patterns (3). These drawbacks limit the application of optogenetics to answer important questions involving manipulation of specific cells such as neurons and other excitable cells like vascular smooth muscle cells and astrocytes, within ensembles. In order to overcome these limitations, it is desirable to achieve optogenetic stimulation with single-cell level precision. One approach is to utilize two-photon absorption, which is characterized by being limited to the immediate vicinity of the focal point, thereby achieving spatially restricted activation of opsin channels (3). Several studies have demonstrated the feasibility of this approach (4-12); however, two-photon optogenetics has relatively low efficiency in eliciting a biological response, which limits its *in vivo* applications (**Figure 1**).

**Figure 1:**
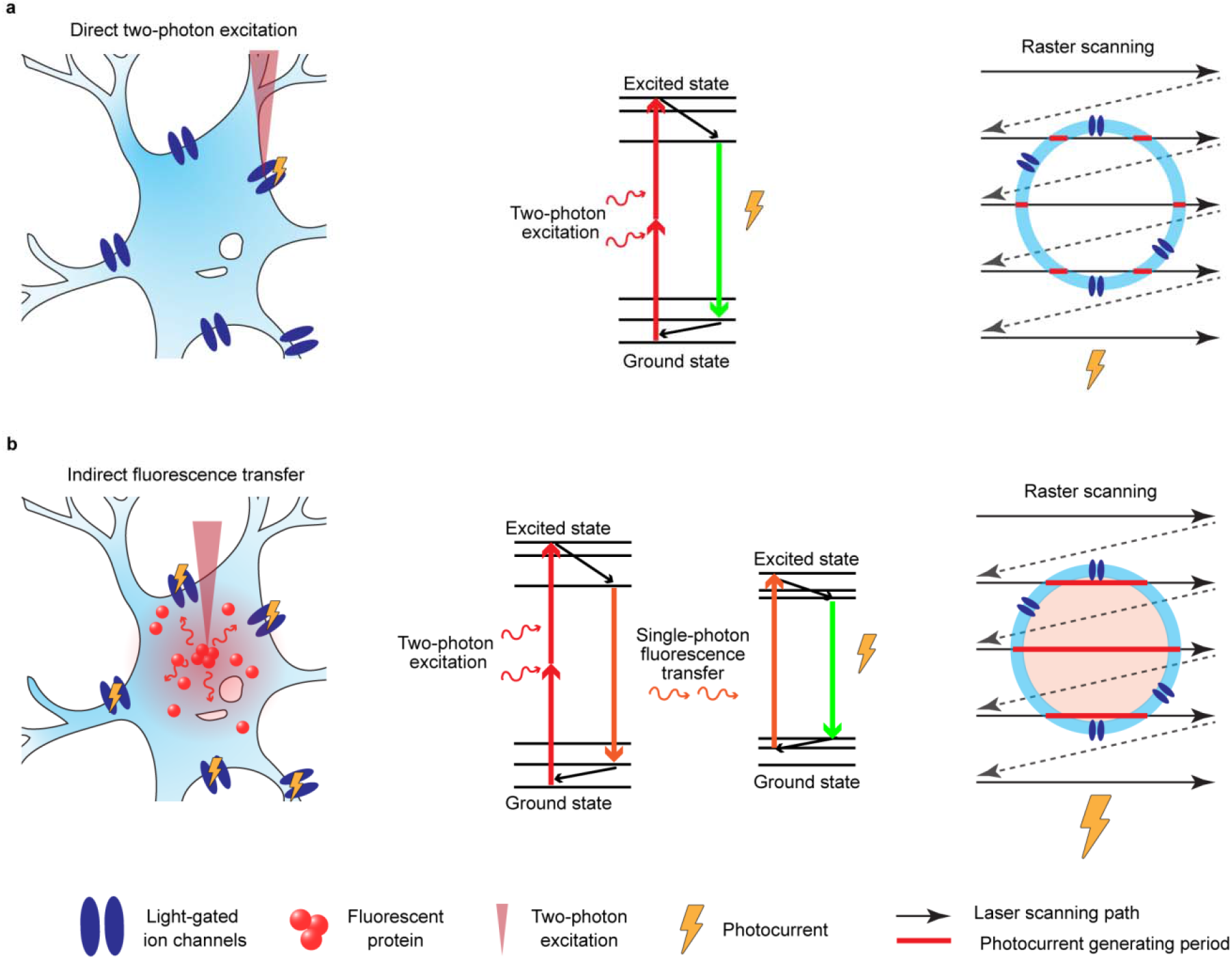
Diagram depicting the principle of two-photon excitation fluorescence transfer (TEFT). (**a**) Direct two photon illumination of light gated ion channels (opsins) induces photocurrents mainly at focal points on the cell membrane as the laser scans the field of view (left diagram). Jablonski diagram depicting standard two photon excitation of opsins and resulting photocurrent (Middle diagram). Raster scanning showing opsin activation at site of focal membrane illumination by two-photon laser (Right diagram). (**b**) Expression of fluorescent proteins or presence of organic dyes in the immediate vicinity of opsins (i.e. cell cytoplasm or intravascular space) allows indirect two-photon illumination (Left diagram) by scanning the entire area and exciting fluorophores all along the path instead of just at sites of opsins on the membrane (Right diagram). The two-photon excitation of adjacent fluorophores generates single photon emissions that are less focal and can indirectly activate the adjacent membrane opsins and generate photocurrents. This can improve the efficiency of opsin activation because it generates a larger number of exciting photons, thereby more efficiently stimulating the adjacent opsins.

One of the reasons for its low efficiency is that the two-photon effect occurs in a submicron focal volume (13). Thus, only a small patch of cell membrane is illuminated at any point in time during laser scanning, which limits the number of opsin molecules that are synchronously stimulated. This limits the ion flux necessary to induce changes in membrane potential and the resulting ability to trigger action potentials. Opsin stimulation can be improved by the use of higher laser power, but unfortunately this can also have direct effects on membrane potential and cell excitability (14-16), likely due to two-photon thermal effects (17, 18), which can cause confounding opsin- or activity-independent ion channel opening. Furthermore, the use of high laser power is problematic as it may induce a variety of cell signaling changes and toxicity (19-21). An alternative to improve the efficiency of two-photon illumination is to use fast laser scanning and generate spiral paths that roughly match the cell’s perimeter, which allows nearly simultaneous activation of a larger number of opsin channels along the cellular membrane (4, 9). A related technique uses spatial light modulators to generate a hologram in the sample so that the laser can simultaneously illuminate the entire target cell (12, 22, 23) and thereby elicit a more robust cellular response. While studies have demonstrated *in vivo* manipulation of neural activity at single-cell resolution with both techniques (24-27), they require advanced optics and complex instrument operation that limit their implementation by most researchers. Thus, it would be of great utility to improve the efficiency by which opsins are stimulated with conventional two-photon illumination and thereby achieve reduced laser scanning times and power requirements. Here we propose a robust and practical approach to achieve *in vivo* optogenetic control of single cells that we termed two-photon excitation fluorescence transfer (TEFT). This approach markedly improves the efficiency of targeted optogenetic control of excitable cells in the live brain and should be entirely compatible with current methodologies including spiral cell scanning and holographic illumination, commonly used for studies of brain networks, cell physiology and neurovascular coupling.

## Results

### Two-photon excitation fluorescence transfer (TEFT) for in vivo optogenetics

In conventional two-photon optogenetics, opsins located in a small patch of membrane defined by the width of the diffraction-limited point-spread-function (PSF) (hundreds of nanometers diameter) are activated at any point in time, leading to relatively inefficient stimulation of the cell (9). Instead of directly activating the light-sensitive opsin channels on the cell membrane, TEFT utilizes the two-photon laser to excite fluorophores located in the vicinity of the opsins. These fluorophores have a fluorescence emission spectrum matching the optimal (single-photon) opsin absorption, which is now used instead of two-photon excitation, to indirectly stimulate opsins in the target cell **(Figure 1)**. Effectively, this converts two-photon stimulation into a local single-photon point source that can be efficiently used for optogenetics (9)(**Figure 2**). In addition, with TEFT, fluorescent molecules can be selected for a high two-photon cross section and high quantum yield that better matches the adjacent opsin excitation properties. Importantly, one can control the concentration of either small fluorophores or fluorescent proteins and their location (i.e. cell cytoplasm or intravascular space) and excite the full volume of the PSF which is much greater than the amount of membrane bound opsins that are normally excited by direct two photon illumination. This translates into a more efficient single-photon illumination point source that can stimulate larger areas of the adjacent cell membranes. While the total amount of fluorescence is determined by the focal excitation, the light intensity that reaches the membrane-bound opsins decays with the inverse square of the distance from the focal point. We further calculated two forms of excitation crosstalk: 1) The out of focus two-photon excitation of a neighbor cell at a different axial position (**Figure 2h**); and 2) The activation of a neighbor cell from the target cell’s fluorescence (**Figure 2i** and **Supplementary Figure 1**). Our calculation shows that the activation of a neighbor cell can only reach up to a maximum of 10% of that of the target cells. This provides a theoretical framework for understanding the ability of TEFT for efficient optogenetic stimulation without a loss of single cell resolution (**Figure 2**).

**Figure 2:**
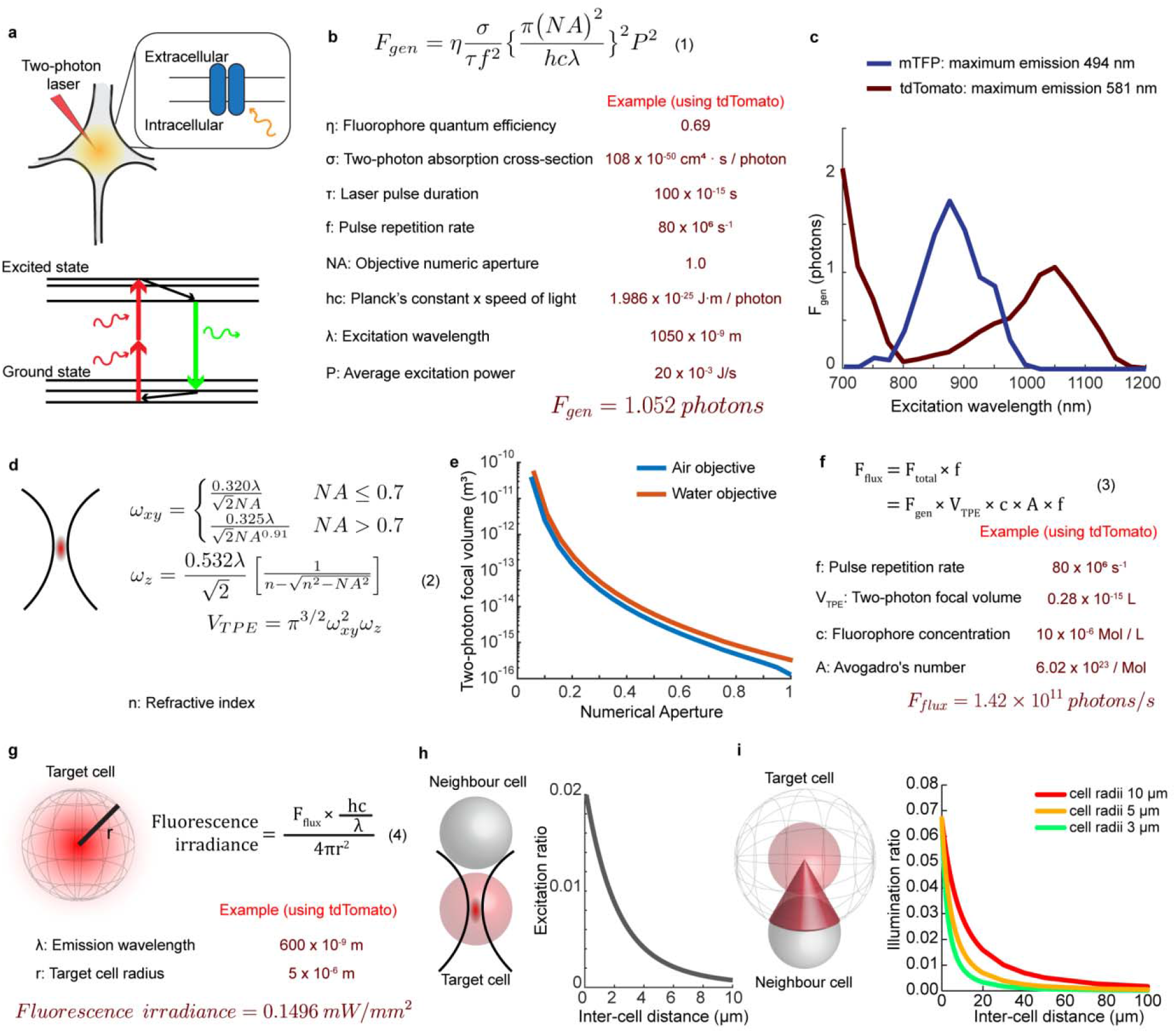
Theoretical estimation of fluorescence irradiance with two-photon excitation. (**a**) Schematic diagram of fluorescence transfer mediated two-photon optogenetics and the Jablonski diagram for the principle of two-photon excitation. Two-photon excitation generates fluorescence that can be absorbed by opsins expressed on the cell membrane. Two prerequisites for efficient fluorescence transfer optogenetics: 1, matching the spectrum between fluorophore emission and opsin absorption; and 2, sufficient irradiance of the fluorescence to generate photocurrents. (**b**) Equations describing the photons generated per fluorophore per pulse (modified from (13)). (**c**) Simulation of two-photon excited fluorescent photon for mTFP and tdTomato at different excitation wavelengths. (**d**) Equations estimating two-photon focal volume (modified from (42)). (**e**) Theoretical calculation of two-photon focal volume at 1050 nm, with air or water objective at different numerical aperture configurations. (**f**) Equations estimating total fluorescence flux. (**g**) Equations estimating fluorescence irradiance to the cell surface. (**h**) Estimation of axial off-target two-photon activation. The plot on the right shows the excitation ratio between the neighbour cell and the target cell at different distances. **(i)** Theoretical estimation of fluorescence transfer efficiency to a non-target cell. The ratio of fluorescence radiation to the target cell (red) and to a non-target neighbor cell (grey) can be expressed as the ratio between the spherical cone cap area to the total sphere area. The plot on the right shows the theoretical calculations of illumination ratios between the target and neighbour cells at different sizes and distances.

### In vivo optogenetic control of vascular smooth muscle cells with TEFT stimulation

Guided by the abovementioned TEFT optogenetics principles, we tested the feasibility of two-photon optogenetic stimulation of vascular smooth muscle cells (vSMC) in the live mouse brain to locally control cerebral blood flow (28). Optogenetic control of the brain vasculature has recently been implemented as a powerful tool for dissecting mechanisms of neurovascular coupling and its control by different vascular mural cell types (29-31). We hypothesized that TEFT may improve the efficiency and reliability of vascular optogenetics, and thus investigated this method with various combinations of opsins and intravascular fluorescent dyes as previously described (32). We first tested the ability to induce vessel constriction in Cspg4-Ai32 mice, in which the perivascular vSMCs express the excitatory ChR2. In order to provide the fluorescence emission that matches the optimal absorption of ChR2, we intravascularly injected cascade blue-conjugated albumin. We then scanned a region of interest (ROI) over the selected vessel segment using the femtosecond laser tuned to 800 nm, a wavelength that is suboptimal to directly excite ChR2 (33) and therefore cannot induce adequate vSMC contraction (**Figure 3**). As predicted, we observed a robust vessel constriction that was only elicited when we implemented the stimulation in the presence of intravascular cascade blue (**Figure 3a, Supplementary Movie 1**). In contrast, we did not observe any vessel constriction when we used an unconjugated control albumin, albumin conjugated with a dye not optimally matched to ChR2 absorption (**Figure 3d**) or when we used a 950-nm wavelength, which does not excite the cascade blue dye (**Figure 3e**). Importantly, the stimulation only induced constriction of the targeted vessel segments, while the diameter of adjacent segments or vessels in the nearby region remained unchanged (**Figure 3g** and **Supplementary Figure 3**). Next, we used archaerhodopsin (Arch)-expressing mice (Cspg4-ArchT (Ai40D)) to induce vSMC hyperpolarization and determine the efficiency of TEFT to induce vSMC-relaxation and consequent vasodilation. To achieve the optimal single-photon activation wavelength of Arch (∼545 nm, (34)), we utilized an intravascular Alexa514-conjugated albumin and 900-nm two-photon illumination. This resulted in efficient and focal vessel dilation (**Supplementary Movie 2**), which did not occur in the absence of the intravascular dye (**Figure 4a and b**). For both ChR2 and ArchT activation, we found that many other dyes with similar emissions were capable of inducing opsin activation. For example, all three blue-emitting fluorescent dyes, Cascade-blue, Alexa 405 and AMCA (aminomethylcoumarin acetate), were able to trigger vessel contraction in Cspg4-ChR2 mice (**Figure 3f**), while the two yellow-emitting dyes, Alexa 514 and Lucifer yellow, produced vessel dilation in Cspg4-ArchT mice (**Figure 4e**). Together, these results demonstrate that the fluorescence generated from these intravascular dyes by two-photon excitation was a potent indirect light source for highly efficient and specific optogenetic control of vSMCs *in vivo*.

**Figure 3:**
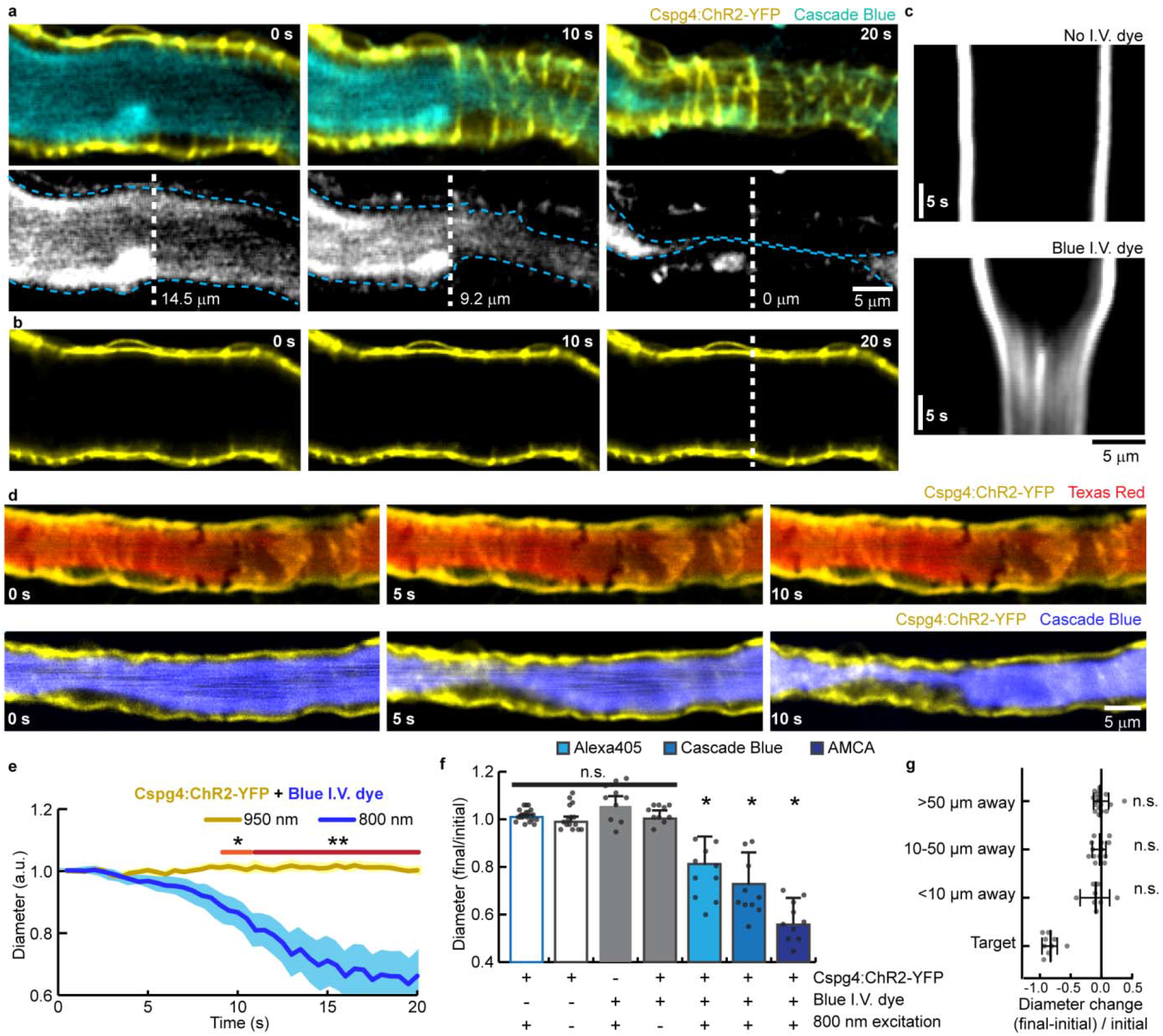
Fluorescence transfer-mediated two-photon optogenetic excitation of vascular smooth muscle cells in vivo. (**a**) Time-lapse intravital brain imaging in mice expressing ChR2 in vascular smooth muscle cells (Cspg4:ChR2-YFP) show focal vessel constriction induced by two-photon illumination of intravascular blue dye (cascade blue). Blue dashed lines (lower row) show the outlines of the intravascular space (cross-section widths indicated by white dashed lines). (**b**) Time-lapse images of the same vessel segment as in **a**, without the intravascular blue dye, showing no changes in diameter with the same laser power. Scanning parameters in **a** and **b**: 25 Hz, 800 nm laser, 10 µs dwell time, 10mW. (**c**) Vessel cross-sections during the scanning periods at the locations of white dashed lines in **a** and **b**. (**d**) Two-photon time-lapse intravital images of optogenetic stimulation in a Cspg4-cre:Ai32 mouse with a red intravascular dye show little change in diameter (upper row), while the same vessel segment with a blue intravascular dye demonstrates robust constriction (lower row). (**e**) Quantification of normalized vessel diameters during two-photon scanning at 800nm and 950nm wavelengths. Data are represented as mean ± standard deviation. N=10 vessels for each group. Orange and red segments indicate statistically significant timepoints between groups (*: p<0.05 and **: p<0.01, respectively, Student’s t-test between groups for each time points, with Bonferroni’s correction for multiple comparisons). (**f**) Quantifications of vessel diameters at the end of the stimulation with different experimental conditions. Data are presented as mean ± standard deviation, with individual datapoints provided (N=10 to 20 vessels per group). One sample Wilcoxon tests were used for each group to compare to 1, with additional Bonferroni’s correction for multiple comparisons (*: p<0.05). (**g**) Quantification of vessel diameter changes between the start and the end of TEFT stimulation in the vessel segments at different distances to the target region. Data are presented as mean ± standard deviation, with individual datapoints provided (N=10 to 20 vessels per group). One sample Wilcoxon tests were used for each group to compare to 0.

**Figure 4:**
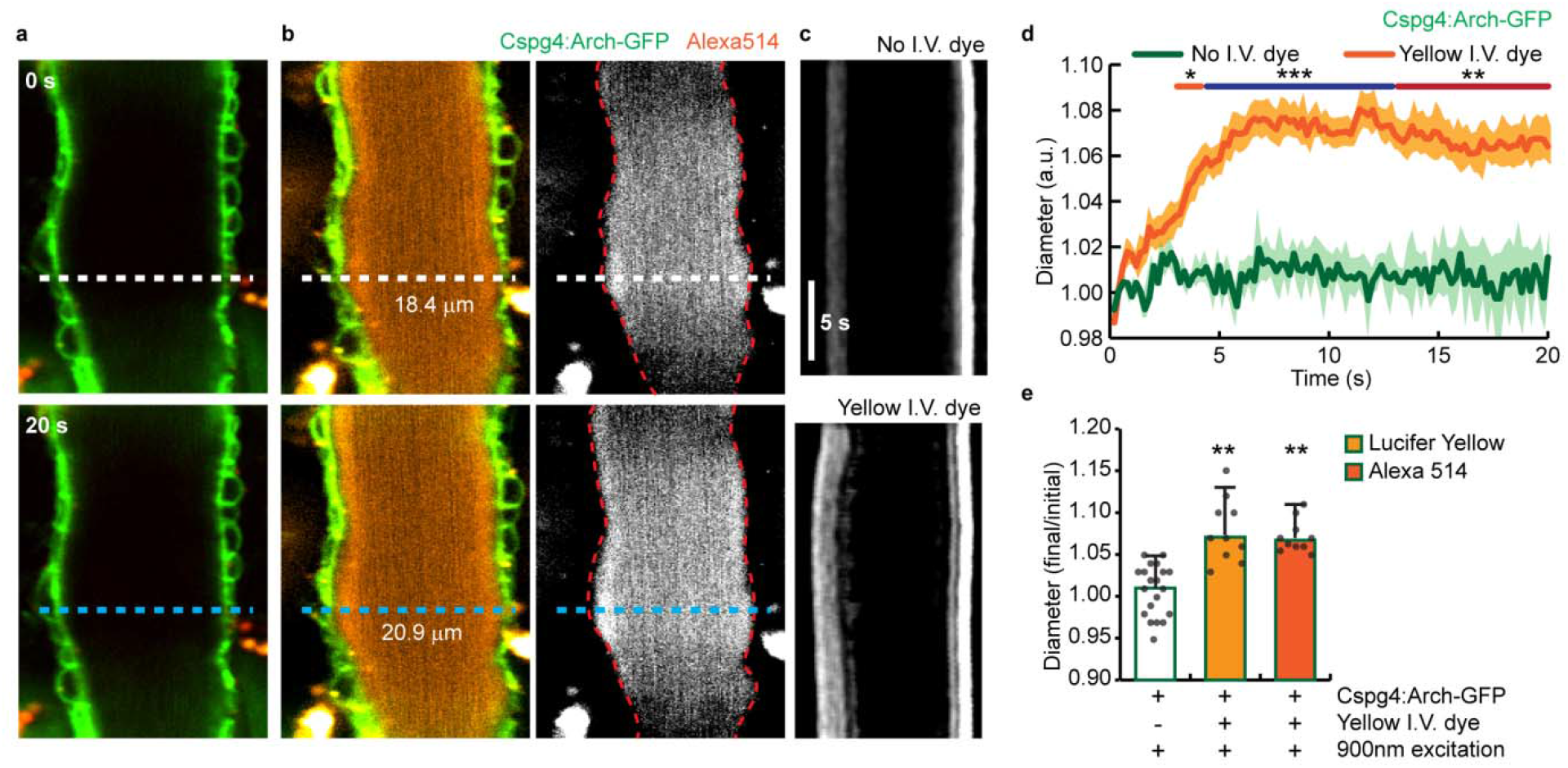
Fluorescence transfer-mediated two-photon optogenetic inhibition of vascular smooth muscle cells in vivo. (**a**) Two-photon time-lapse images of vessel dilation in archaerhodopsin expressing mouse (Cspg4:ArchT-GFP). (**b**) Representative two-photon time-lapse images of the same vessel segment with yellow intravascular dye. White and blue dashed lines show site where diameter was measure overtime in **a** and **b**. Scanning parameters in **a** and **b**: 25 Hz, 900 nm laser, 10 µs dwell time, 10mW. (**c**) Vessel cross-section line profiles depicted overtime during scanning at the locations of dashed lines in **a** and **b**. (**d**) Quantification of normalized vessel diameters with and without yellow intravascular dye. Data are represented as mean ± standard deviation. N=10 vessels per group. Yellow, red and blue segments indicate statistically significant timepoints between groups (*: p<0.05; **: p<0.01 and ***: p<0.001, respectively, Student’s t-test between groups for each time point, with Bonferroni’s correction for multiple comparison). (**e**) Quantification of vessel diameters with different experimental conditions. Data are presented as mean ± standard deviation, with individual datapoints provided. N=10 to 20 vessels per group. One sample Wilcoxon tests were used for each group to compare to 1, with additional Bonferroni’s correction for multiple comparison (**: p<0.01).

### Improved efficiency of two-photon optogenetics in neurons by TEFT

Having demonstrated the effectiveness of TEFT using intravascular dyes to activate vSMCs, we next explored the feasibility of applying this method to neurons. In order to measure the efficiency of optogenetics, we performed patch-clamp recordings of pyramidal neurons in acute mouse brain slices (**Figure 5**). We implemented TEFT, by expressing ReaChR in neurons (through *in utero* electroporation) and filling the recording electrode with Alexa594 solution (**Figure 5a**). To achieve optogenetic stimulation, we scanned these cells by two-photon illumination of a ROI covering the entire cell body using a wavelength of 800 nm for ∼50 ms. We found that adding the dye increased the peak current by ∼50%, compared to cells with no dye, under the same laser stimulation conditions (**Figure 5b and c**). In addition, TEFT-mediated activation showed faster onset kinetics compared to direct two-photon optogenetics (**Figure 5d**), consistent with the TEFT principle, that all opsins on the target cell were activated simultaneously, rather than sequentially as is the case with direct two-photon membrane-opsin activation.

**Figure 5:**
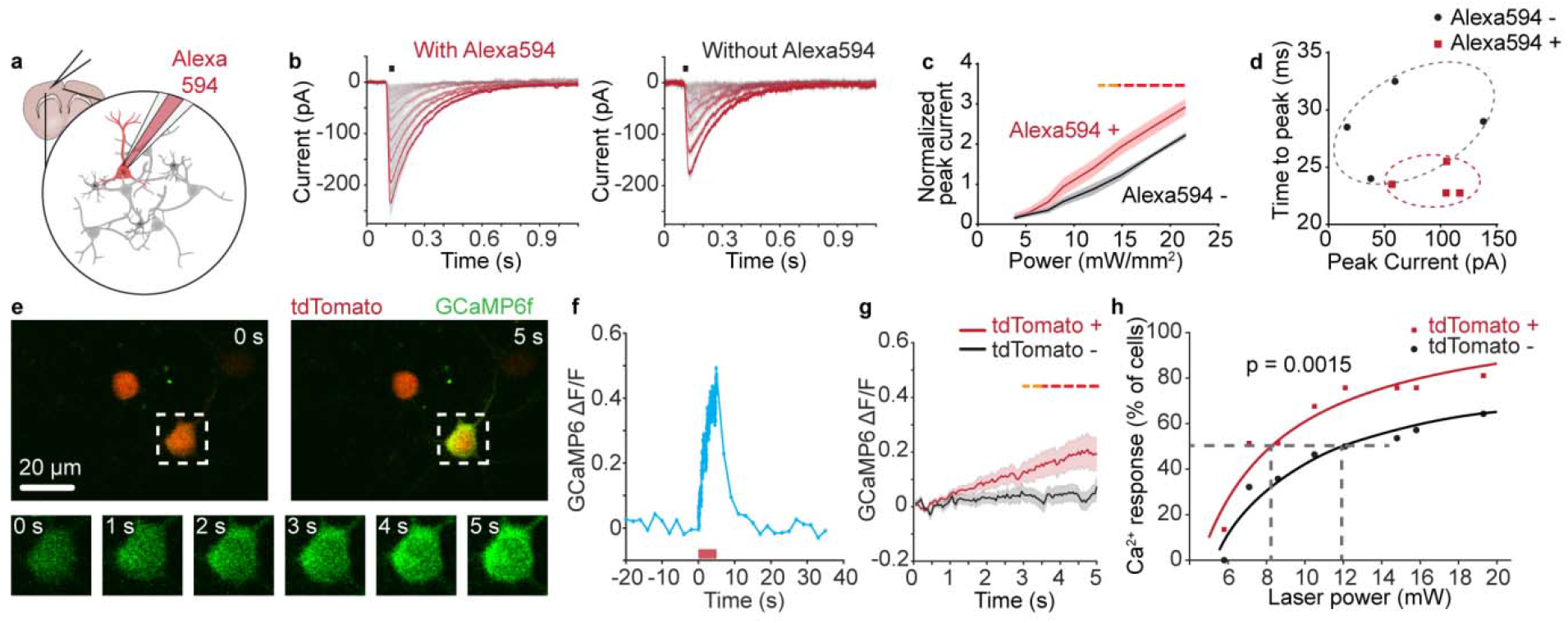
In vivo optogenetics of single neurons using fluorescence transfer-mediated two-photon stimulation. (**a**) Schematics of patch clamp recording with TEFT. (**b**) Representative traces of two-photon elicited currents in cells with or without Alexa594 dye at 800 nm. Laser powers are 3.8, 5.4, 7.3, 8.9, 12, 14.8, 18.1 and 21.6 mW, for curves from grey color to red. Black lines indicate a 50 ms two-photon stimulation period. (**c**) Quantification of the peak current amplitudes in cells with or without Alexa594 dye, normalized by the peak current recorded at 920 nm at 5 mW. N = 4 cells in each group. Data are represented as mean ± standard error (Student’s t-test comparison between groups for each time point, a false discovery rate of 5% was set for multiple comparison correction. Yellow dash lines: p<0.05, red dash lines: p<0.001) (**d**) Quantification of onset phase kinetics of two-photon activated currents in cells with or without Alexa594 dye, plotted against peak currents. Dash lines indicate the perimeter for each condition group. (**e**) Two-photon raster scanning in a live mouse brain of ROI (white dashed square) covering a neuron that is co-expressing ReaChR, GCaMP6 and tdTomato, induces robust calcium transients (ROI scan parameters: pixel size: 0.42 µm/pixel, 50 Hz, 5 s, 920 nm laser, 4 µs dwell time, 8.6 mW). Time-lapse images (bottom panel, green) show rapid increase in calcium levels following two-photon illumination. (**f**) GCaMP6 calcium response of a neuron coexpressing ReaChR, GCaMP6 and tdTomato using the same scanning parameters as in **e** (ROI scanning interval indicated by the orange bar). (**g**) Comparison of two-photon optogenetics in ReaChR/GCaMP6 positive neurons with and without tdTomato expression at 8.6 mW excitation power (N=37 for tdTomato positive; N=28 for tdTomato negative). Data are represented as mean ± standard error (Student’s t-test comparison between groups for each time point, a false discovery rate of 5% was set for multiple comparison correction. Yellow dash lines: p<0.05, red dash lines: p<0.001). (**h**) GCaMP6 responses by 920nm two-photon activation of ReaChR in cells with and without the expression of tdTomato. Curve fitting showed that tdTomato-expressing neurons are more efficiently activated than tdTomato-negative neurons at various laser powers (20% rise of GCaMP6 fluorescence used as arbitrary threshold of neuronal activation) (p=0.0015 comparing differences between fitted curves, see methods for statistical details).

We next applied TEFT in the live brain. Due to the difficulty of introducing organic fluorescent dyes into cells *in vivo*, we instead overexpressed fluorescent proteins in the target neurons. We used *in utero* electroporation of neurons in the mouse brain to first co-express the red-emitting fluorescent protein tdTomato and the opsin ReaChR (peak absorption 595 nm), as well as the calcium sensor GCaMP6 to detect neuronal activity changes. To test the feasibility of implementing TEFT in neurons, we scanned these cells by two-photon illumination of a ROI covering the entire cell body using a wavelength of 920 nm, which has been reported to excite ReaChR (35). This focused scanning triggered a rapid rise in GCaMP6 fluorescence (**Figure 5e and f**).

We then compared this optogenetic-induced calcium rise in cells with and without tdTomato co-expression. While two-photon scanning can directly stimulate ReaChR at relatively high powers (35), co-expression with tdTomato markedly increased the efficiency, and enabled activation with laser powers that are normally too low for ReaChR stimulation (**Figure 5g**). We observed ∼50% reduction of the laser power required to reach 50% probability of activation using the TEFT method (**Figure 5h**). Importantly, TEFT improves the efficiency of cell activation while maintaining the single-cell resolution, since we could elicit a robust calcium rise in the targeted cell, without any calcium changes in the immediately adjacent one (**Supplementary Figure 3a-c**). Together, these observations demonstrate that TEFT-mediated two-photon optogenetics improves the efficiency of cell activation.

### Fluorophore/opsin constraints critical for optimal TEFT efficiency

In contrast to the tdTomato/ReaChR pair for TEFT optogenetic stimulation, we were not able to elicit two-photon activation of ChR2 when pairing it with genetically encoded blue fluorescent proteins or SNAPtag-targeted organic dyes co-expressed in the same neurons (**Supplementary Figure 4**). This contrasts with the highly efficient activation we observed when ChR2 in vSMCs was stimulated in the presence of blue intravascular organic dyes (**Figure 2**). As an explanation for this phenomenon, we hypothesized that the number of photons emitted by the donor fluorophores in the vicinity of opsins is a critical variable that determines their efficient activation. To better understand this relationship, we calculated the theoretical number of photons emitted after two-photon excitation of various well-known fluorescent proteins (36, 37)(**Figure 2 and Supplementary Figure 4c**). With these data, we determined that the mTagBFP/ChR2 pair that we used experimentally for neurons, was not suitable for TEFT optogenetics, given that for laser powers of ∼10 mW, typical of most intravital applications, the calculated emitted fluorescence of mTagBFP was only of the order of 0.01 mW/mm^2^, which is two orders of magnitude lower than the reported power needed for ChR2 activation (38). In contrast, with the tdTomato/ReaChR pair, using 10 mW for two-photon illumination, yielded around 0.15 mW/mm^2^ (**Supplementary Figure 4d**), which is known to be sufficient to elicit strong ReaChR photocurrents (39). One way to overcome the low two-photon cross section of most genetically encoded blue fluorescent proteins, would be to increase their intracellular concentration to achieve greater net photon emissions. However, it is difficult to increase their intracellular concentrations beyond ∼10 micromolar (40). This contrasts with the concentration of intravascular dyes that we used for stimulation of ChR2 in vSMCs (∼500 micromolar), which can be further increased as needed, thereby achieving highly efficient TEFT optogenetic stimulation.

## Discussion

Here we reported a novel approach to improve the limited efficiency of two-photon illumination for opsin stimulation *in vivo*. By positioning organic dyes or genetically encoded fluorescent proteins in the cytoplasm or immediate vicinity of opsins (intravascular), and using two-photon illumination to excite them, a focal source of single-photon emissions is generated, which efficiently activates adjacent opsins. The TEFT technique retains the focal illumination properties (given the rapid intensity decay as a function of distance from the single-photon light source, see **Figure 2**), which allows opsin stimulation at cellular and possibly subcellular resolution. Interestingly, our calculation showed that TEFT has a low (<7%) cross-talk illumination power to the neighbor cell, across different sizes of the target cell (**Figure2** and **Supp. Figure 1**). Together with the fact that the effective illumination power is higher in smaller structures (inversely correlated with surface area, **Figure 2**), TEFT may provide an ideal method for optogenetic experiments requiring high resolution such as those of single dendritic spines. We demonstrated that TEFT improves the efficiency of *in vivo* experiments otherwise not easily achievable such as targeted opsin stimulation of vSMCs and neurons with widely available standard two-photon microscopy setups. The lower laser power requirements achieved by this method could be critical for reducing thermal injury (41) and unwanted laser-induced electrophysiological effects independent of opsin activation (14, 16). TEFT can be further optimized in the future by improving the quantum yield of the paired fluorescent proteins utilized or by developing more efficient methods for targeting bright organic dyes to specific cellular compartments, thereby achieving higher *in vivo* concentrations. Finally, this method is entirely compatible and should also improve the efficiency of other methods for two-photon optogenetic stimulation such as the use of fast spiral scanning paths (4, 9) or scanless holographic approaches (22, 23). Together our data demonstrates a significant improvement in the methodologies for targeted cell optogenetics stimulation that are critical for experiments requiring precise spatial and temporal single-cell stimulation for investigation of cellular physiology and neural networks *in vivo*.

## Figures and figure legends

**Supplementary Movie 1: Fluorescence transfer-mediated two-photon optogenetic activation of ChR2 in vascular smooth muscle cells leads to vessel constriction**. Time-lapse videos of the same vessel segment in a Cspg4-Ai32 mouse with (right panel, 800nm excitation) and without (left panel, 925nm excitation) the excitation of the intravascular blue-emitting dye. Scanning parameters in both: 25 Hz, 10 µs dwell time, 10mW laser power.

**Supplementary Movie 2: Fluorescence transfer-mediated two-photon optogenetic activation of ArchT in vascular smooth muscle cells leads to vessel dilation**. Time-lapse videos of the same vessel segment in a Cspg4-Ai40D mouse with (right panel, 900nm excitation) and without (left panel, 930nm excitation) the excitation of the intravascular yellow-emitting dye. Scanning parameters in both: 25 Hz, 10 µs dwell time, 10mW laser power.

## Materials and Methods

### Mice

All rodent procedures were approved by the Yale University Institutional Animal Care and Use Committee. For vascular studies, transgenic mice that express the Cre recombinase under the mural cell NG2 (Cspg4) promoter, and reporter lines with cre-dependent channelrhodopsin-2 (Ai32) or Archaerhodopsin-3 (Ai40D) were purchased from The Jackson Laboratory (JAX# 008533, JAX# 021188, JAX# 012569). Cre-expressing strains were crossbred with the reporter strains and the offspring were used for all experiments. For neuronal studies, wild type mice were used for electroporation of various constructs (JAX# 000651). For all experiments, 2-3-month-old mice from both sexes were used.

### Reagents

Purified albumin (Sigma-Aldrich, 05470) was used for fluorescent dye conjugation. Reactive esters were used for labeling (Thermo Fisher Scientific, C-2284, A6118, A30000, L-1338) according to the manufacturer’s instruction. The labeled albumin was diluted so that 5mg reactive dyes constituted 1mL of injection stock. 100µl of labeled albumin was injected intravenously before imaging, final dye concentration in blood was estimated to be ∼0.5mM. In all conjugations, albumin was used at concentrations greater than the number of fluorophores to eliminate the need for free fluorophore purification.

To express constructs by in Utero electroporation we obtained and modified the following plasmid constructs from Addgene: CAG-tdTomato, CAG-ReaChR (#50954), Syn-GCaMP6f (#100837), CAG-tagBFP (#49151), Syn-ChR2 (#58880), CAG-jRCaMP (#61562). See **Supplement** for maps of modified plasmids.

### In utero electroporation

In utero electroporation was done as previously described (43). Briefly, Plasmids were used at the final concentration of 1.0 μg/μl (for each plasmid), mixed with 2 mg/ml Fast Green for visualization during plasmid injection and electroporation. Electroporation was performed around embryonic day 13 to 15 (E13 to E15). Mice were anesthetized with ketamine/xylazine (100mg/kg and 10mg/kg i.p.). Buprenorphine was administered (i.p.) every 12 hours for 2 more days following surgery. After exposing the uterine horns, ∼1 μl of plasmid mixture was pressure injected into the lateral ventricle of each embryo via a pulled glass microelectrode (tip size 10∼20 um) using Picospritzer II (General Valve, 20 psi). 50 V current pulses generated by a BTX 8300 pulse generator (BTX Harvard Apparatus) were used for electroporation. Mice were allowed to age to 1 month prior to utilization in all experiments.

In order to verify the co-expression of tdTomato, ReaChR and GCaMP6, we used in vivo two-photon imaging to monitor the cells while shining a red LED to trigger ReaChR activation, triple positive cells should have baseline red fluorescent signal and a LED-triggered green fluorescent signal. The co-expression probability was very high with electroporation. And only cells that demonstrated robust triple expression were then targeted for two-photon stimulation.

### Craniotomy surgery, window implantation and in vivo two-photon imaging

Mice were anesthetized with an intraperitoneal injection of Ketamine/Xylazine mixture, with final concentration of 100mg/kg and 10mg/kg, respectively. The status of anesthesia was assessed periodically with hind paw pinch. The mouse was head-fixed to a custom-made headplate by gluing the skull to it. A craniotomy of about 4mm diameter was made (AP -1.5mm, ML 2.0 mm) with a dental drill, with dura mater carefully removed. A coverslip was put to cover the craniotomy opening and secured with cyanoacrylate glue. The mouse was kept anesthetized during subsequent imaging sessions, and immediately euthanized after finalizing the experiment.

Two-photon imaging was carried out with a commercial system (Bruker Ultima Investigator), controlled through Prairie View software. A tunable Ti:Sapphire laser was used to generate two-photon excitation with its wavelength and mode-locking tuned through MaiTai software. In the case of RCaMP imaging, a 1045nm fixed wavelength laser (MaiTai InSight X3) was used. A pockels cell was used to modulate laser power; and the laser power on the sample was measured with a power meter (Thorlabs PM100D). The point scanning was achieved by galvanometer scanners with various dwell times. The full frame rate was kept at 0.5 Hz, and for stimulation, the scanning within regions of interests (ROIs) was at 20 Hz or 50 Hz frequencies. During ROI scanning, the regions outside the ROI were not scanned nor imaged (represented by the dashed portion of the GCaMP6 response curved in **Figure 5**). Fluorescence emission was collected with gallium arsenide phosphide photo-multiplier tubes. A 20x water immersion 1.0 numerical aperture objective (Zeiss) and a 10x air 0.4 numerical aperture objective (Leica) were used for most experiments. For all in-vivo experiments, imaged blood-vessels were identified within 250 μm from pia mater, and imaged neurons were located at depth between 100 to 350 μm. In all our experiments, cells demonstrated normal spontaneous and stimulated responses, and showed no signs of persistently elevated calcium levels associated with cell damage (44).

### Single cell patch clamp and two-photon optogenetics

Acute brain slices of the in utero electroporated mice (P30-P40) were prepared following a N-methyl-D-glucamine (NMDG) protective recovery method (45). Whole cell patch clamp and two-photon optogenetics were then performed in slices in an ASCF containing (in mM) 120 NaCl, 3.1 KCl, 1.1 CaCl2, 1.2 MgCl2, 1.25 MgSO4,26 NaHCO3, 0.5 L-glutamine, 0.1 ascorbic acid, 0.1 Na-pyruvate, and 20 glucose; saturated with 95% O2–5% CO2 at 35°C. To target fluorescent cells, we used a two-photon microscope system (Ultima; Prairie Technologies) equipped with a Ti:Sapphire pulsed laser (MaiTai), configured on an Olympus upright microscope (BX51WI) with a 20×, 0.5 NA objective lens (LUMPlanFL/IR) and a 60x, 1.0 NA objective lens(LUMPLANFL/IR). Cells were patched under 60x objective lens with pipette solutions as follows (in mM): (1) for voltage clamp, 105 CsMeSO4, 0.5 CaCl2,10 HEPES, 5 EGTA, 5 Na2-phosphocreatine, 2 ATP-Mg, 0.5 GTP-2Na, 2 ascorbic acid, and 8 QX314-Cl (pH 7.2), with 20–30 CsOH; (2) for current clamp, 105 potassium gluconate, 5 KCl, 0.5 CaCl2, 2 MgCl2, 5 EGTA, 10 HEPES, 5 Na2-phosphocreatine, 2 ATP-2Na, 0.5 GTP-2Na, and 2 ascorbic acid (pH 7.2) with 5 NaOH and 15 KOH. Liquid junction potential was calculated with pCLAMP software and corrected (Molecular Devices, Union City, CA). Once whole cell patch clamp was achieved, 20x objective lens was switched for two-photon optogenetics. A single ROI (∼10 × 10 µm) including only the cell soma was chosen for raster scan, with 4 µs dwell time, and laser intensity less than 25 mW (3.8, 5.4, 7.3, 8.9, 12, 14.8, 18.1 and 21.6 mW), which shows no clear photo damage to cell membranes. 800 nm wavelength was used to excite intracellular dye (Alexa 594, 0.5 mM). The photocurrent under 920 nm, which would not activate Alexa 594(46) was also recorded to evaluate the expression level of ReaChR. An external voltage was used to trigger the two-photon image scan, so that the timing of laser scanning and cell voltage/current can be accurately matched for later analysis. The raster scanning was started at 0.1 s in each trail and last for 50 ms.

### Statistics

Statistical analyses were carried out using GraphPad Prism (8.4.1). Data were presented in mean ± standard deviation in **Figure 3**, and in mean ± standard error in **Supplementary Figure 4**. For comparing normalized vessel diameter time-lapse traces (**Figure 3e** and **4d**), Student’s t-test was performed with each timepoint, with Bonferroni correction for multiple comparison. For comparing the two-photon mediated vessel motility in different conditions, one-sample Wilcoxon tests were used for each group to compare to a value of 1 (**Figure 3f** and **4e**) or 0 (**Figure 3g**), with additional Bonferroni’s correction for multiple comparison. For comparing the efficacy of neuronal optogenetics with and without fluorescence transfer (**Supplementary Figure 4e**), we fit the response probability from each group to the following exponential equation: Y = 1-exp(-K*(X-L), in which the parameter L indicates the minimal power to elicit calcium responses and K indicates the change rate of the curve. Extra sum-of-squares F test was used to determine whether two sets of parameters were statistically different.

### Data availability

All data presented show individual data points whenever possible to facilitate replication of our graphs and associated statistics. All raw images and recording data are available upon request.

### Code availability

All equations for modeling calculations were presented in figures. All data processing steps in the study used standard built-in functions that should be self-explanatory. The scripts used for analyzing calcium responses are included in **Supplementary Material**. We will provide any additional details of the functions upon request.

## Acknowledgement

We thank Dr. Oscar Hernandez (Stanford University) for his guidance on the estimation of fluorescence irradiance by two-photon excitation, and Dr. Jonathan Demb (Yale University) for helpful conceptual and technical discussions. This work was supported by NIH R01NS115544 (J.G.). J.B. was additionally supported by NIH R01GM118486. M.C. and Z.J.Z were additionally supported by NIH R01EY026065 (ZJZ), P30EY026878 (Yale Vision Core). P.Y. was additionally supported by startup fund IDF2641002 (Fudan University).

## Author contributions

P.Y and J.G. designed the study. L.T., P.Y. and F.C. carried out the experiments. P.Y. and J.B. performed theoretical calculations. Y.X., M.C. and J. Z. performed the electrophysiology recording. P.Y., L.T. and J.G. wrote and edited the manuscript.

## Supplementary Material

In some cases, the DNA sequences encoding the target proteins (tagBFP, tdTomato and ReaChR) were cloned from the original constructs into an AAV-CAG backbone (the backbone was acquired from Addgene #28014, deleting the GFP sequence), and the resulted plasmids were used for in utero electroporation. Standard molecular cloning procedures were performed using the following reagents: Phusion High-Fidelity DNA Polymerase (New England BioLabs, M0530S), Restriction enzymes (New England BioLabs), T4 DNA ligase (ThermoFisher Scientific, EL0011), QIAGEN plasmid Maxi kit (#12162). All modified constructs were sequences to verify the correct insertion and sequence. The modified plasmid maps are showed below:

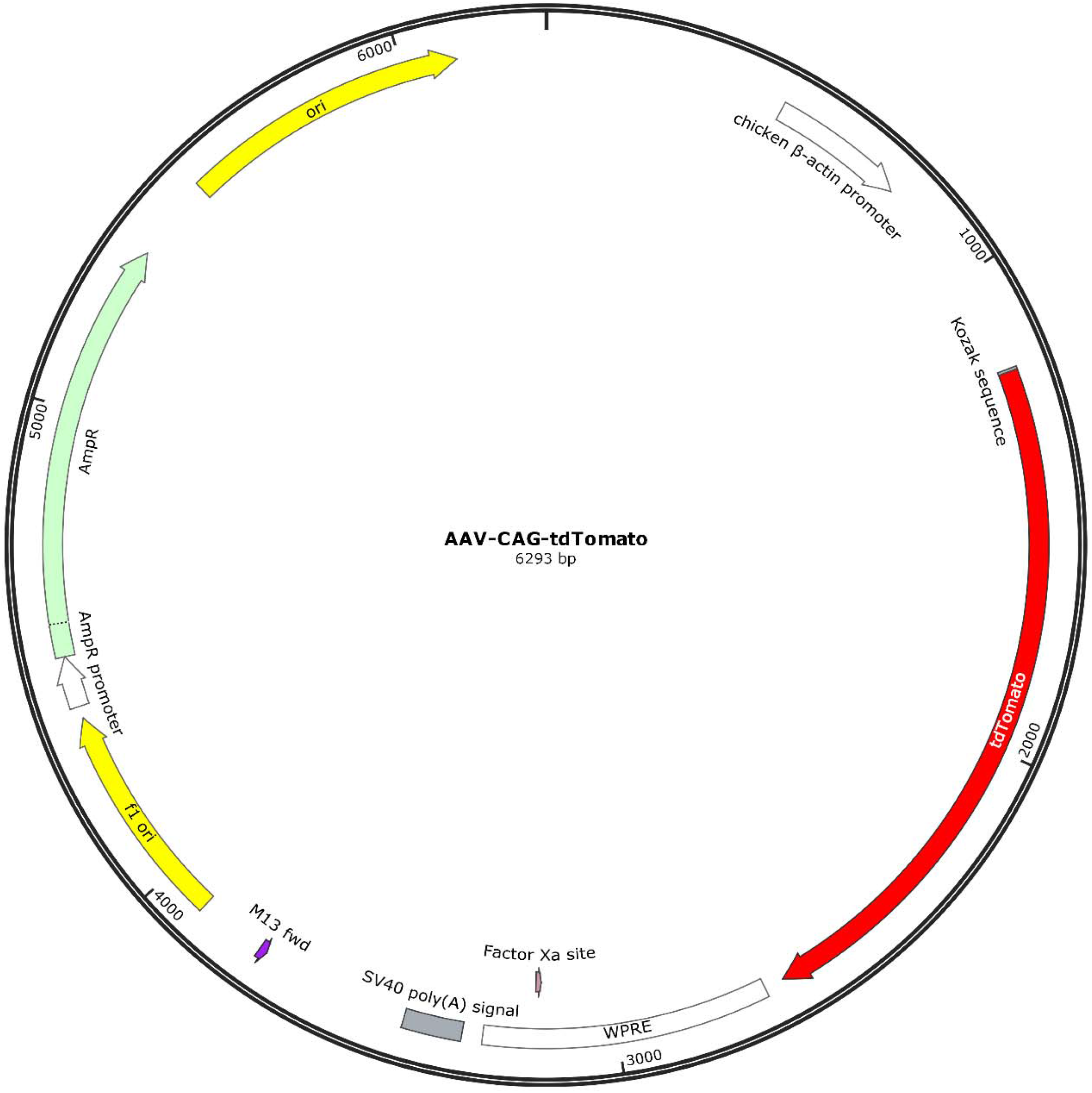

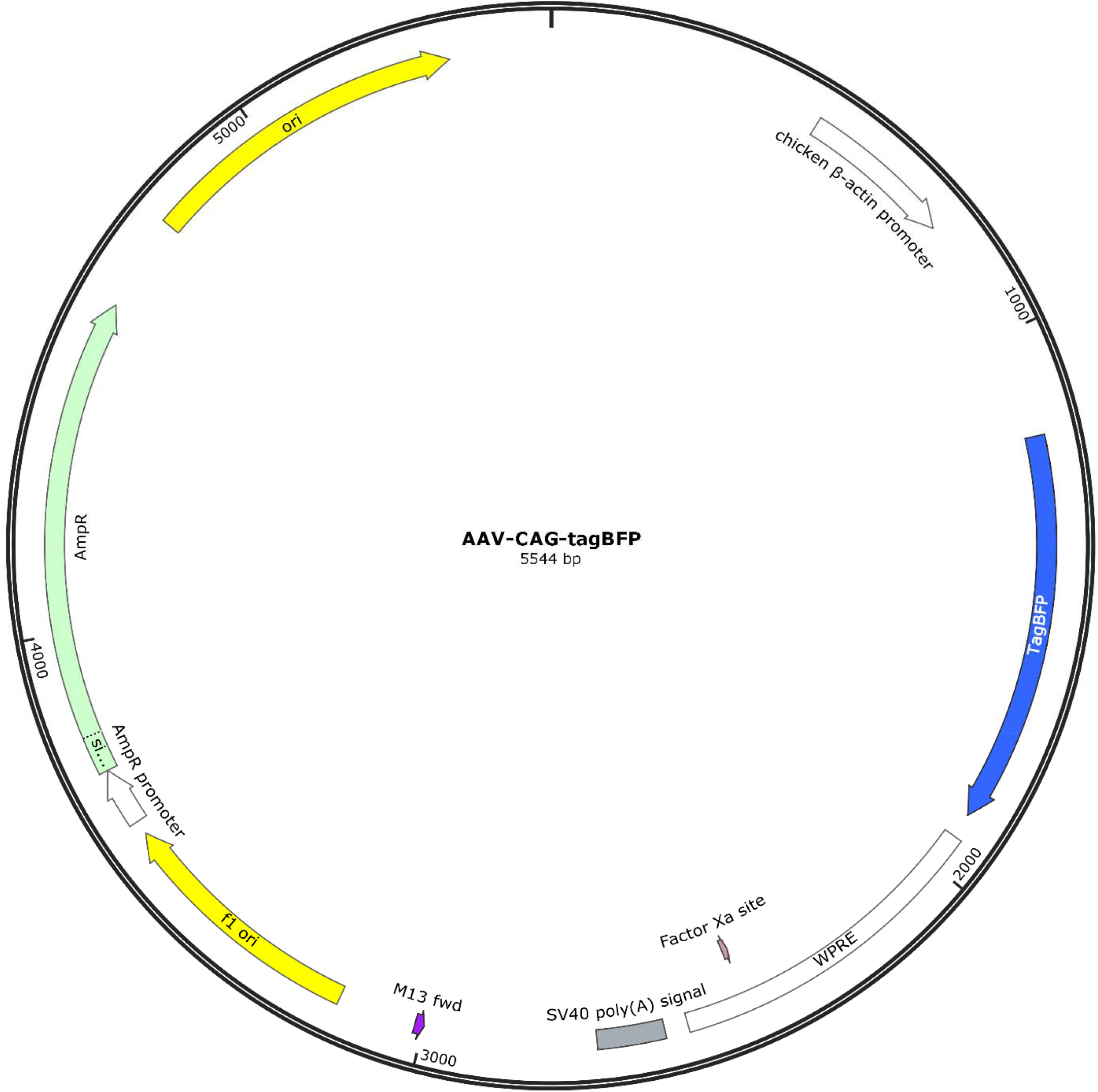

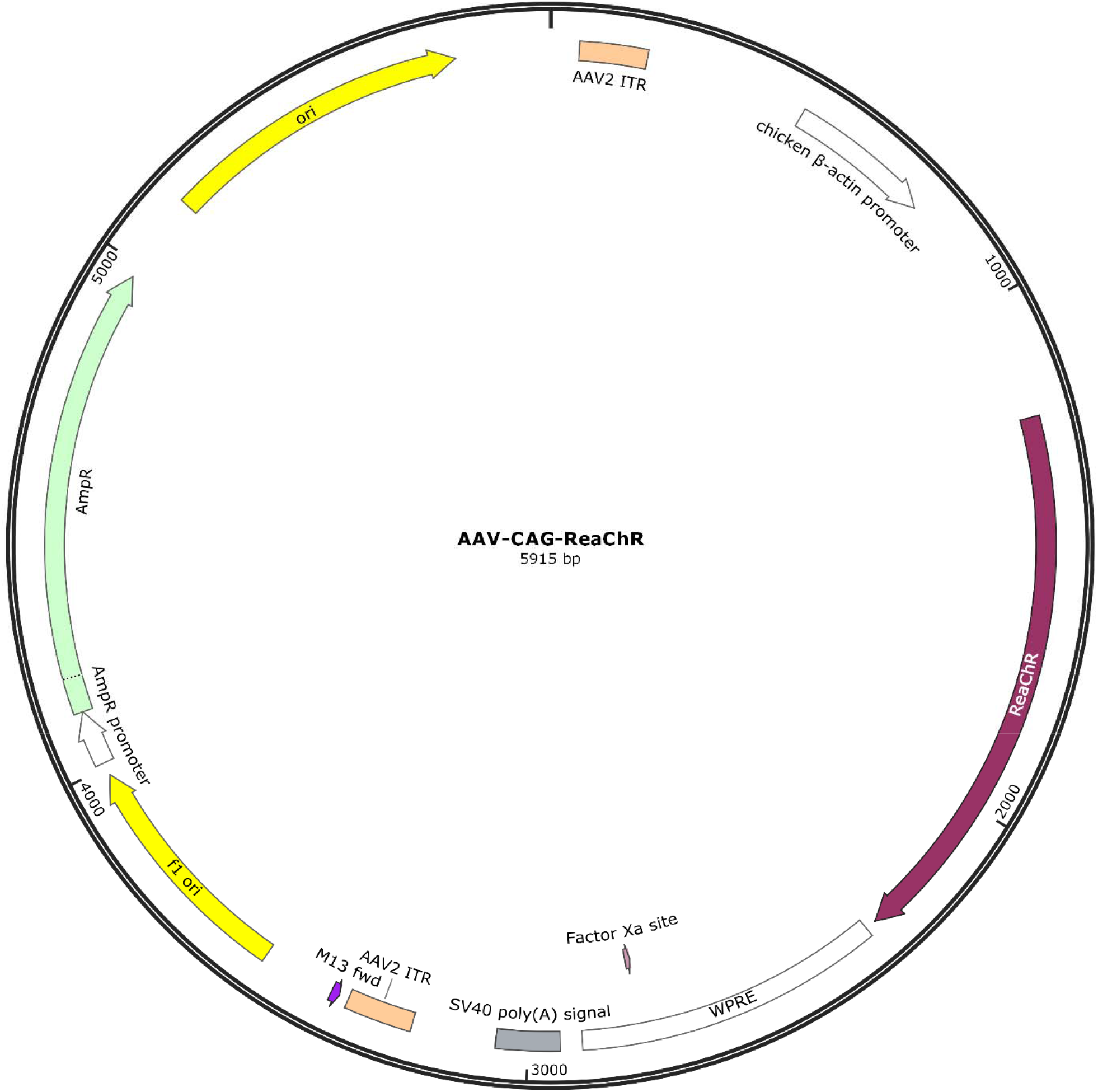

## Custom scripts for analyzing calcium responses

We used MATLAB for this part of the analysis.

~~~
%%
clearvars; close all;
[title,path] = uigetfile(‘*.xml’,’Select the image file’);
cd(path);
text = fileread(title);
% Find dimensions of 5D image
% XY dimensions: Entries assumed to be the same for pixel dimensions
xdim = str2double(regexp(text, …
 ‘PVStateValue key=“pixelsPerLine”.*?value=“(\d*)”‘, ‘tokens’,’once’));
ydim = str2double(regexp(text, …
 ‘PVStateValue key=“linesPerFrame”.*?value=“(\d*)”‘, ‘tokens’,’once’));
% Z axis dimension
z_cell = regexp(text, ‘Frame relative.*?index=“(\d*)”‘, ‘tokens’);
zslice = cellfun(@(x) str2double(x{1}),z_cell);
% T axis dimension
t_cell = regexp(text, ‘Sequence type=.*?cycle=“(\d*)”‘, ‘tokens’);
tpoints = cellfun(@(x) str2double(x{1}),t_cell);
if tpoints == 0; tpoints=1; end
t = cellfun(@str2double, regexp(text, …
 ‘Frame relativeTime=“(\d+(\.\d*)|0)”‘, ‘tokens’,’all’))’;
% Channel dimension (img names found on same line, parsed also)
ch_img_names_cell = regexp(text, …
 ‘File channel=“(\d*)”.*?filename=“(.*?)”‘, ‘tokens’);
ch = cellfun(@(x) str2double(x{1}),ch_img_names_cell);
img_names = cellfun(@(x) x{2},ch_img_names_cell, ‘UniformOutput’, 0);
% Bit depth of img in filesystem tif_bit_depth = str2double(regexp(text, …
 ‘<PVStateValue key=“bitDepth”.*?value=“(\d*)”‘, ‘tokens’,’once’));
% Create 1×1 mapping between img_name, channel, z slice, timepoint
% Channel index
ch_ind = ch;
% Zslice and timpoint index need to be repeated to match img_names elements
flat = @(x) x(:);
z_ind = flat(repmat(zslice, [numel(unique(ch_ind)) 1]));
t_ind = flat(repmat(tpoints, [numel(unique(z_ind))*numel(unique(ch_ind)) 1]));
% Initialize image specified datatype
xyczt_img = zeros(ydim, xdim, max(ch), …
 max(zslice), max(tpoints), ‘uint16’);
% Read individual tif files into 5D img
for n = 1:numel(img_names)
 xyczt_img(:,:,ch_ind(n), z_ind(n), t_ind(n)) = imread(img_names{n})…
 * double(intmax(‘uint16’)/2^tif_bit_depth);
end
clearvars -except xyczt_img t;
temp = xyczt_img(:,:,:,:,2);
xyczt_img = temp;
%%
% ROI of the neuron, BFP run this section
bw = imbinarize(imgaussfilt(mat2gray(xyczt_img(:,:,4,1)),2),0.4);
figure,imshowpair(mat2gray(xyczt_img(:,:,4,1)),bw.*0.5);
bw = repmat(bw,[1 1 size(xyczt_img,4)]);
%% ROI of the neuron, tdTomato run this section
bw = imbinarize(imgaussfilt(mat2gray(xyczt_img(:,:,2,1)),2),0.2);
figure,imshowpair(mat2gray(xyczt_img(:,:,4,1)),bw.*0.5);
bw = repmat(bw,[1 1 size(xyczt_img,4)]);
%%
rcamp = squeeze(xyczt_img(:,:,2,:));
intensity = mean(reshape(rcamp(bw),[],size(xyczt_img,4)));
gfp = squeeze(xyczt_img(:,:,4,:));
intensity_gfp = mean(reshape(gfp(bw),[],size(xyczt_img,4)));
intensity = intensity./intensity_gfp.*mean(intensity_gfp);
intensity_m = intensity’./mean(intensity(1:10))-1;
figure,plot(intensity_m);
num2clip(intensity’);
%%
[xData, yData] = prepareCurveData(t(1:150), intensity_m(1:150));
ft = fittype(‘poly1’);
[fitresult, gof] = fit(xData, yData, ft);
p = coeffvalues(fitresult);p(1)
~~~

## Supplementary Figures

**Supplementary Figure 1:**
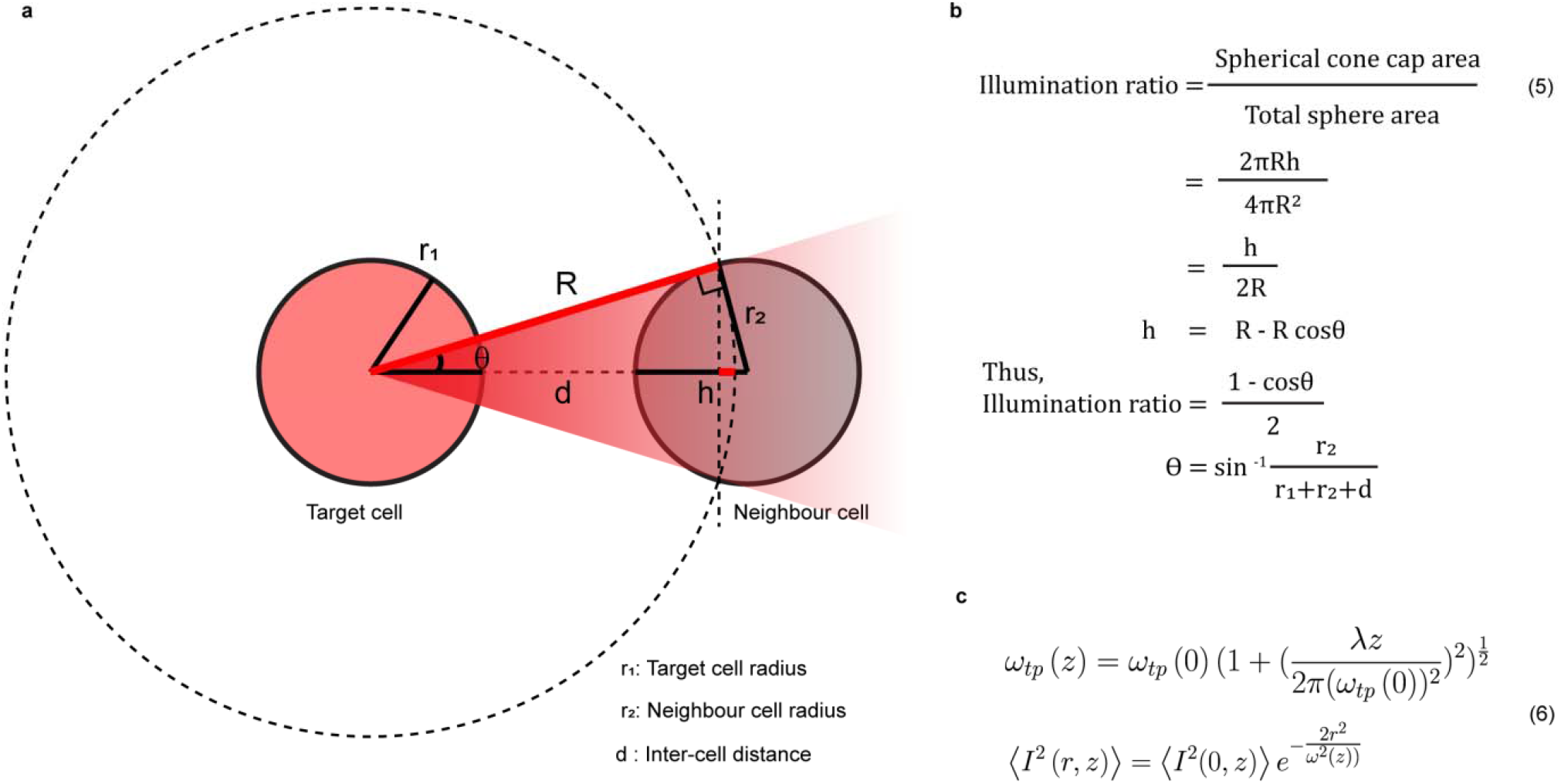
Estimation of the TEFT illumination to a neighboring cell. (**a**) A 2-dimensional schematic illustrating the illumination to the target cell and to the neighbor cell. The total flux through any spherical surface enclosing the fluorescence emission volume is the same and can be considered 100% of the spherical surfaces, while only a cone of the original spherical emission sphere will irradiate a neighboring cell (red cone). (**b**) Equations for calculating the illumination ratio between the target cell and the neighbor cell. The illumination ratio equals to the ratio between the spherical cone cap area and the total sphere area. Results are presented in **Figure 2i**. (**c**) Equations for calculating the out-of-focus two-photon effect along the axial direction for **Figure 2h**.

**Supplementary Figure 2:**
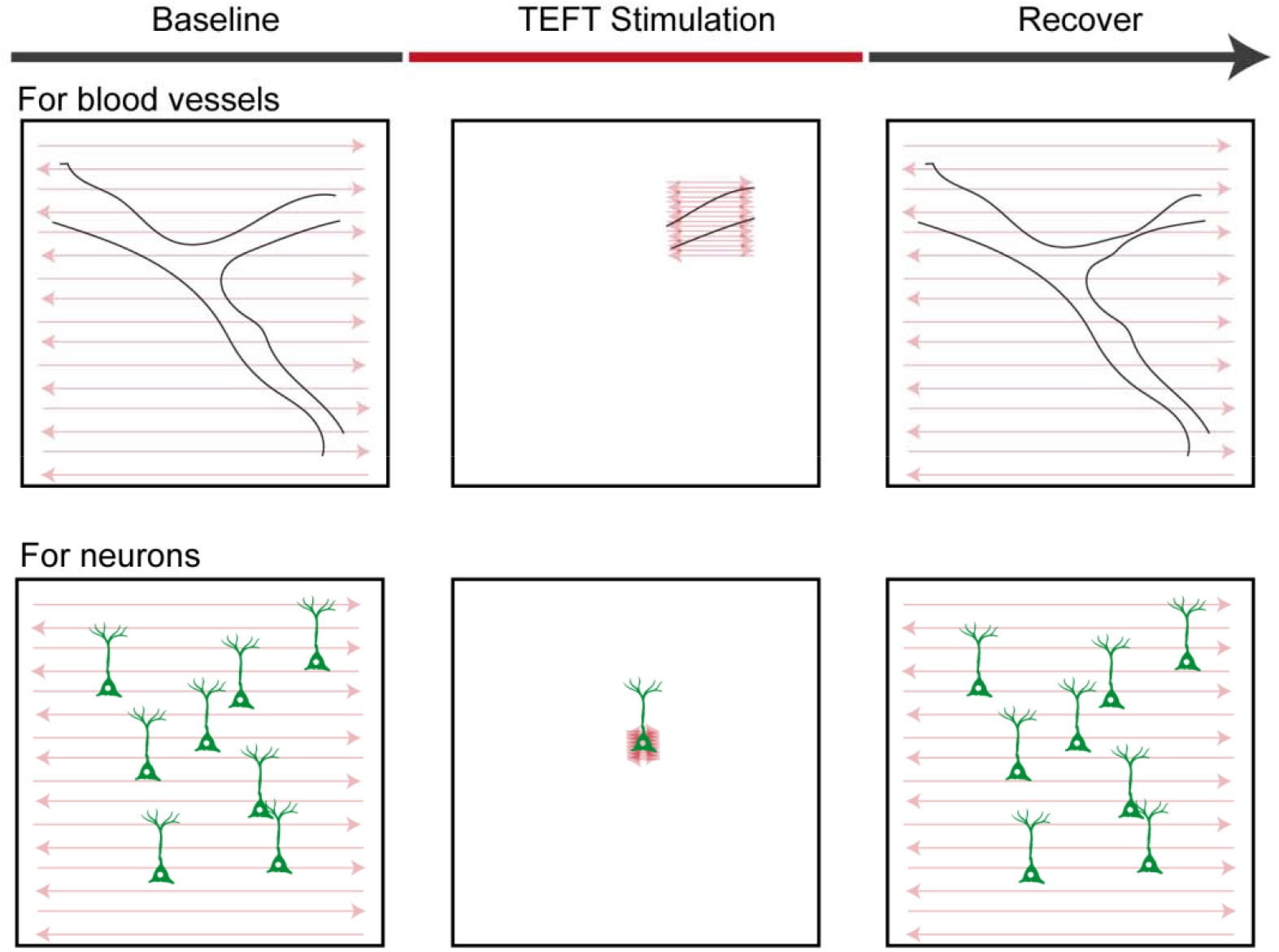
Schematics of the TEFT-imaging protocol used in this study. We implemented a 3-stage imaging protocol to achieve recording and TEFT stimulation. At baseline, the galvanometer was set to full range with low laser power (controlled by a pockel cell). This allowed us to image the target and the surrounding regions without causing activation of the opsin. To achieve TEFT stimulation, we limited the scanning to the location of the target structure with the laser power required for stimulation. We were able to image the target structure during this period. At the end of the stimulation period, the scanning and laser were set back to the same conditions as the baseline stage, and at this point we went on to record the morphology and activity of the same region after stimulation.

**Supplementary Figure 3:**
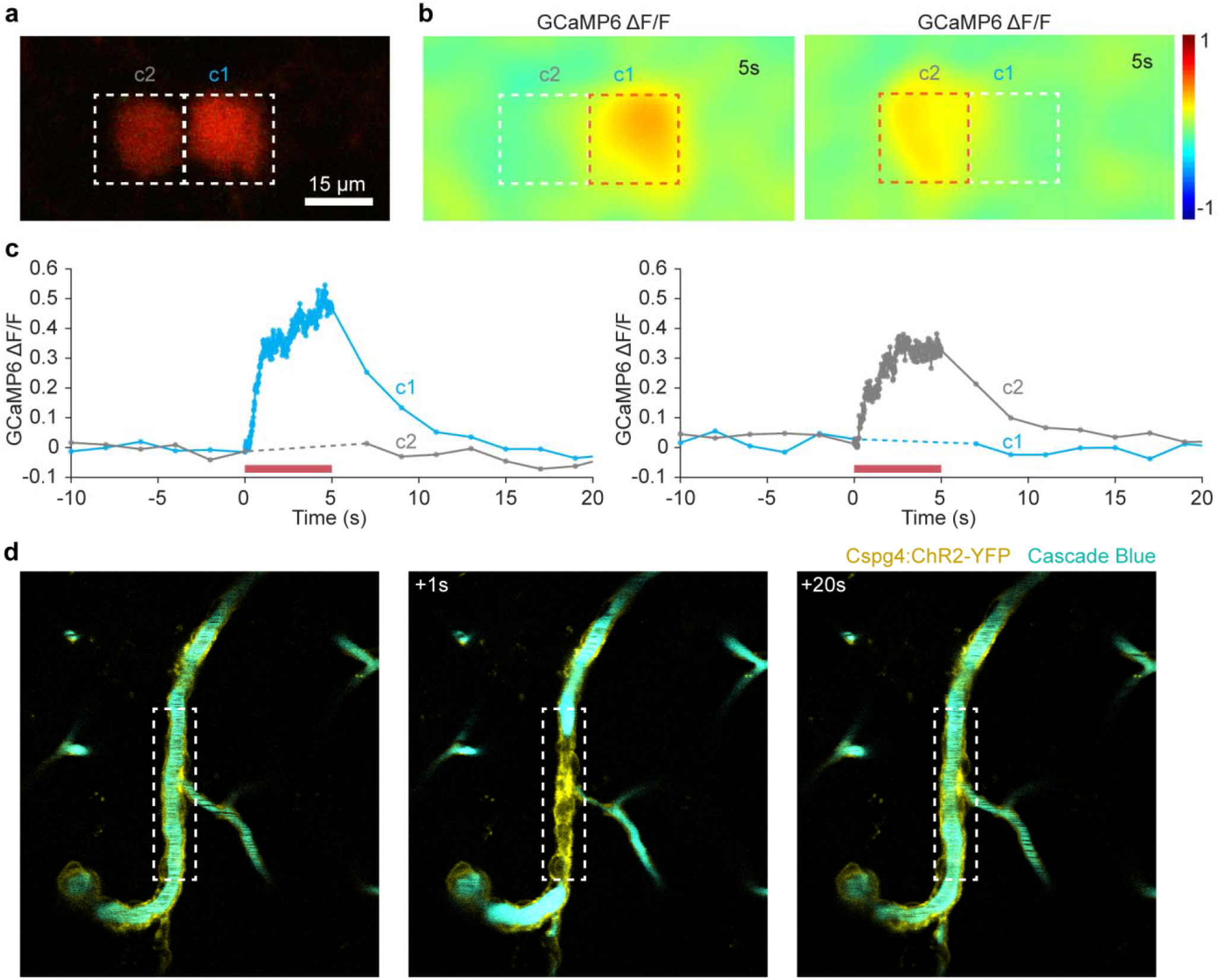
Fluorescence transfer-mediated two-photon stimulation retains high spatial resolution in vivo. (**a**) Sequential two photon stimulation (920nm) of single neuron out of two adjacent ReaChR, GCaMP6 and tdTomato coexpressing cells (top-left). (**b**) Heat maps of individual cell GCaMP6 responses immediately after excitation of a ROI positioned on the stimulated neuron (orange dashed box; middle and -right panels). (**c**) Calcium response curves of these two neurons (Dotted line indicates interval without scanning, see methods for detail) using stimulation parameters as in **a**, show that despite their proximity, the tdTomato emission within the illuminated ROI from either cell is unable to activate the immediately adjacent cell, highlighting the preservation of spatial specificity when using two photon florescence transfer optogenetics. (**d**) Two-photon time-lapse images of vessel constriction in a Cspg4-cre:Ai32 mouse with intravascular blue dye. White dashed boxes indicated the stimulation scanning region of interest. Notice the restriction of the constricted area to the illumination ROI only. Quantification of these data are presented in **Figure 3g**.

**Supplementary Figure 4:**
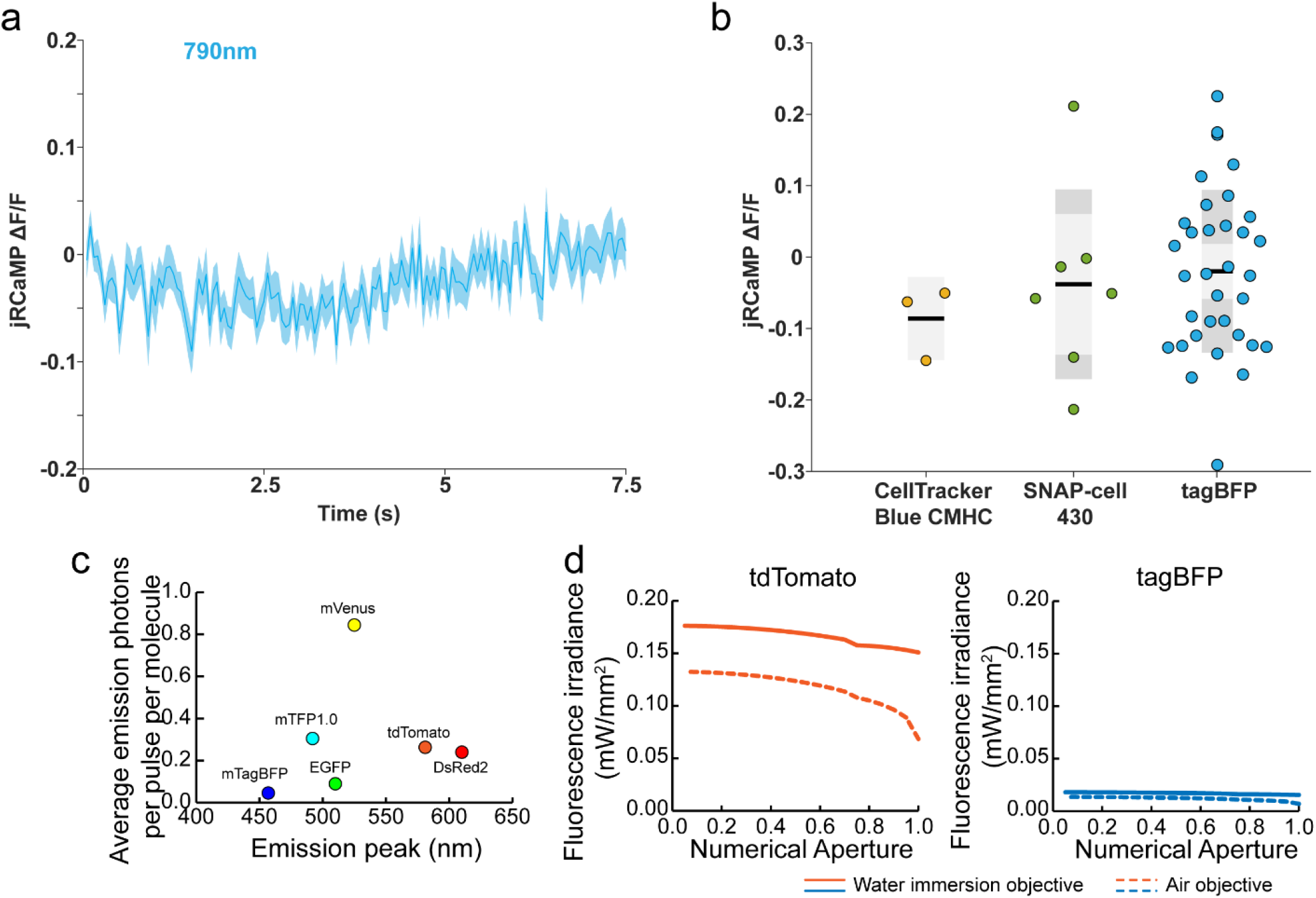
Ineffective excitation of ChR2 in neurons using blue fluorescent proteins reveal critical constraints for optimal TEFT efficiency. (**a**) Calcium response of ChR2, jRCaMP and tagBFP positive neurons (N=34) shows no response when scanned with two-photon (780 nm for tagBFP excitation, 1045 nm for jRCaMP calcium imaging, 4 µs dwell time, ∼15 mW); ROI scanning (48×48 pixels, 20 Hz, for 5 s) performed on the neuron soma. Data are represented as mean ± standard error. (**b**) Quantification of calcium response during two-photon scanning (similar to **a**) with CellTracker Blue CMHC, SNAP-cell 430, and tagBFP in ChR2 and jRCaMP co-expressing neurons (N=3, N=7, N=34). Data are represented as mean ± standard error. An important note is that none of the organic dyes or SNAP-cell 430 labeling strategies achieved sufficiently bright labeling in vivo, potentially limiting the overall stimulation efficiency. (**c**) Theoretical calculations of two-photon fluorescence photon generation for selected fluorescent proteins using the equation in **Figure 2**. Two-photon excitation wavelengths used in calculations corresponded to the peak cross-section of each protein. Laser power was kept at 10 mW with a 1.0 numerical aperture lens. (**d**) Estimations of two-photon excitation fluorescence irradiance of tdTomato and tagBFP, with various objectives (air or water immersion, numerical aperture from 0.05-1.0). Laser power was kept at 10 mW, fluorescent protein concentration was set at 10 μM, and a target cell was 5 μm in diameter. Based on these calculations the blue-emitting proteins have a theoretical low efficiency for exciting the opsins, consistent with our experimental data.

## Supplementary Discussion

We would like to provide an additional discussion regarding whether the red fluorescent protein commonly fused with opsins would be a potent TEFT source, as this question has been brought up multiple times from several readers.

It is important to recognize that the 1:1 ratio between opsin and fluorphore and their location on the membrane both limit the generation of sufficient photons. In fact, the dense packing of fluorescent proteins is a key factor for the efficiency of the TEFT method.

To illustrate this point, here we described a back-of-envelope calculation to put this into proper scale. The key issue is that a fusion protein cannot provide enough photons to induce the TEFT effect.

<u>For TEFT illumination</u>, we used the parameters described in **Figure 2f**:

Two-photon focal volume: 0.28 × 10^−15^ L, fluorophore concentration: 10 μM, this leads to a total of 1686 molecules stimulated at each pulse of two-photon excitation.

Number of molecules = focal volume x concentration x Avogadro’s number

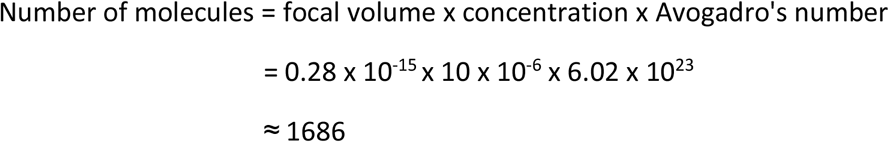

<u>For membrane fusion</u>, the expression of opsins can be ranging from 90 - 750 molecules per μm^2 1,2^. Given the ∼0.2 μm XY-plane resolution, the activated molecules on the membrane are fewer than 30, which is about 2 orders of magnitude lower than the TEFT methods. In fact, the estimated total density of membrane proteins is about 30000 molecules per μm^2 1^, and usually the overexpression of exogenous proteins will be ∼2-3 % of this value. Even if we are assuming very generously that the opsin expression constitutes 10% of total membrane protein, which is never achieved, we are still only activating ∼120 molecules at each moment, which is one order of magnitude lower than our TEFT method.

